# Flexible early prospection of potential behavior in working memory

**DOI:** 10.1101/2023.03.01.530584

**Authors:** Rose Nasrawi, Sage E.P. Boettcher, Freek van Ede

**Author notes:** Correspondence: Rose Nasrawi, Freek van Ede.

## Abstract

For visual working memory to serve upcoming behavior, it is crucial that we prepare for the potential use of working-memory contents ahead of time. Recent studies have demonstrated how the prospection and planning for an upcoming manual action starts early after visual encoding, and occurs alongside visual retention. Here, we address whether such ‘output planning’ in visual working memory flexibly adapts to different visual-motor mappings, and occurs even when an upcoming action will only potentially become relevant for behavior. Participants performed a visual-motor working memory task in which they remembered one or two visual items for later (potential) use. We tracked planning of upcoming behavior through contralateral attenuation of beta-band activity – a canonical motor-cortical EEG signature of manual-action planning. This revealed how action encoding and subsequent planning alongside visual working memory (1) reflects anticipated task demands rather than specific visual-motor mappings, (2) occurs even for actions that will only potentially become relevant for behavior, and (3) is associated with better performance for the encoded item, at the expense of performance to other working-memory content. This reveals how the potential prospective use of visual working memory content is flexibly planned early on, with consequences for later performance.

## INTRODUCTION

Working memory allows us to retain past visual information to guide and prepare for potential future courses of action. For example, when driving a car, you may sequentially check your surroundings through your mirrors, and check your navigation system for the route, before shifting your gaze back to the road. You can use working memory representations of the surrounding traffic and your route to anticipate and prepare for what you might do next: overtake the slow truck in front of you, reduce your speed, or take the next exit. From the perspective that working memory is ultimately aimed at prospecting and guiding potential future behavior (Chatham & Badre, 2015; Fuster & Bressler, 2012; Myers et al., 2017; van Ede & Nobre, 2023; Vries et al., 2020), a central question is when and how this preparation for future actions takes place alongside the encoding and retention of visual information in working memory.

Information in working memory often becomes relevant for guiding behavior several seconds after sensory encoding. Accordingly, future action plans may form gradually during the memory delay (cf., Rainer et al., 1999), similar to the type of action planning observed preceding voluntary movement (Pfurtscheller & Berghold, 1989; Stancák & Pfurtscheller, 1996), or alongside perceptual decision-making (Donner et al., 2009; Lange et al., 2013). By contrast, a recent study (Boettcher et al., 2021) revealed how the consideration of an anticipated future action commenced early after visual encoding. This pattern of *action encoding* – or “output planning at the input stage” – occurred even in the face of an intervening task and predicted performance several seconds later (Boettcher et al., 2021). We build upon this finding, and upon complementary recent findings demonstrating manual action planning alongside visual working memory (Ester & Weese, 2022; Henderson et al., 2022; Nasrawi & van Ede, 2022; Rösner et al., 2022; Schneider et al., 2017; van Ede et al., 2019). Specifically, we address three important questions left unaddressed by previous research.

First, aforementioned studies linking visual representations to manual actions have done so by associating specific visual features to the required response hand. We and others have done so specifically by linking left- and rightward visual tilt (from a vertical reference) respectively to responses with the left- and right hand (Boettcher et al., 2021; van Ede et al., 2019). If action encoding truly reflects the consideration of prospective task demands, it should be invariant to specific visual-motor mappings. In the current study, we therefore counterbalanced the tilt-response mapping to isolate the flexible prospection of future actions alongside visual working memory, independent of visual tilt.

Second, in complex everyday environments (such as our car-driving example above), we often need to hold multiple visual representations in working memory, that serve multiple potential future courses of action (Cisek & Kalaska, 2010; Ester & Weese, 2022; Nasrawi & van Ede, 2022; van Ede et al., 2019; van Ede & Nobre, 2022). Yet, to date, the described pattern of action encoding in working memory has only been demonstrated when just one visual stimulus was relevant for a single upcoming action that was certain to become relevant (Boettcher et al., 2021; see also: Schneider et al., 2017). In the current study, we address whether the early prospection of a future memory-guided action also occurs after the encoding of visual stimuli that will *potentially* become relevant for future behavior.

Finally, in Boettcher et al. (2021) stronger encoding of a certain future memory-guided action was associated with better task performance. By here considering a situation with multiple potentially relevant memory items, we can address whether stronger encoding of a potential future action is generally associated with better performance. Alternatively, potential-action encoding might give rise to a performance trade-off whereby stronger encoding of one potential future action benefits the associated memory representation, at the expense of another.

We show that: (i) action encoding truly reflects the early prospection of a potential future task, adapting flexibly to specific visual-motor mappings; (ii) both certain and potential future actions are prospected immediately after the encoding of visual information into working memory; and (iii) stronger encoding of a potential action facilitates performance of the corresponding visual representation, but this comes at the expense of another.

## RESULTS

Participants performed a visual-motor working memory task (**Figure 1**), in which they were asked to memorize the orientation of colored bars, and reproduce the orientation of one of these bars after a delay using a response dial. We build upon a previously developed and used task (Boettcher et al., 2021; van Ede et al., 2019) with three key manipulations.

**Figure 1.**
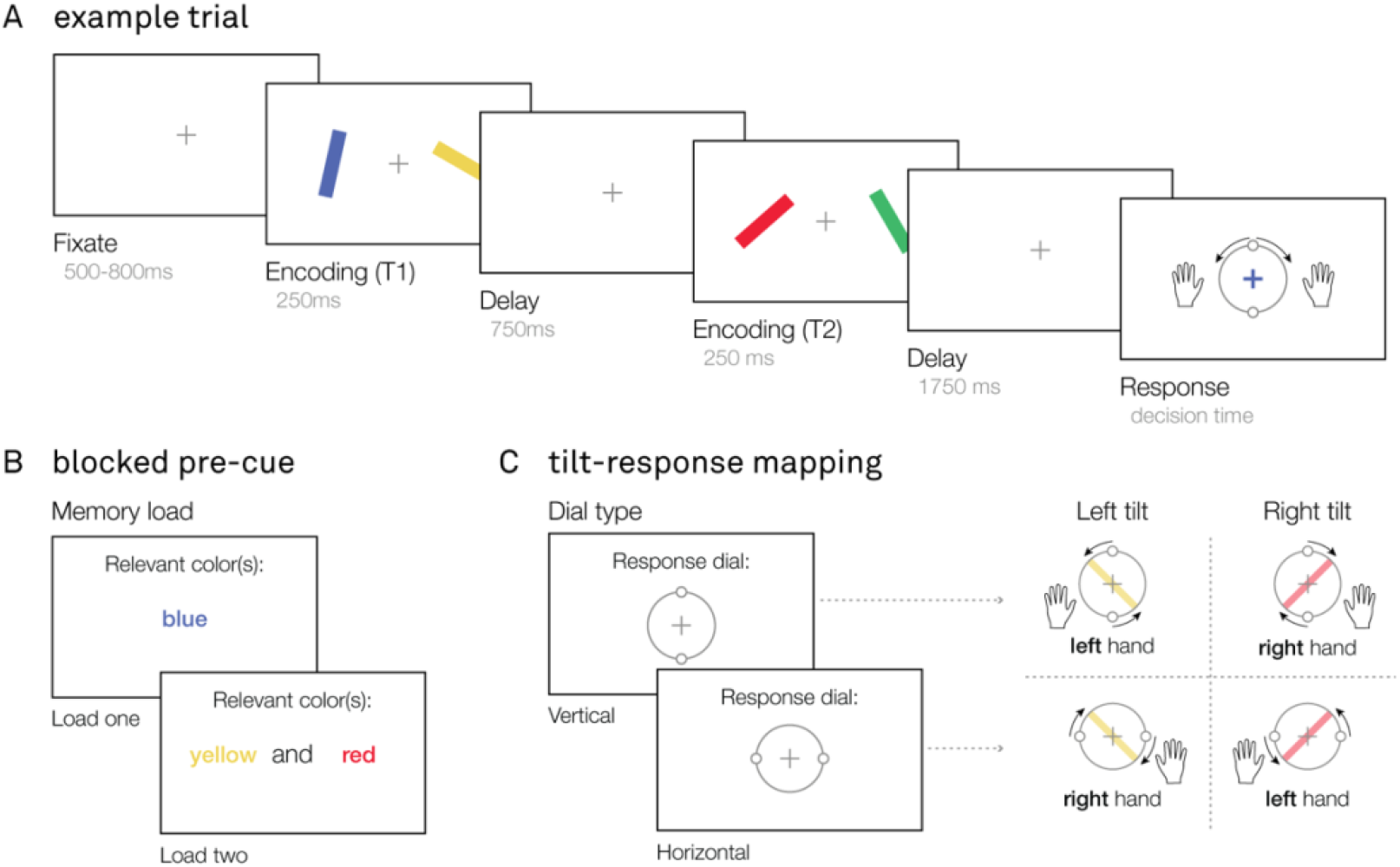
Visual-motor working memory task. (**A**) Example trial. After a brief fixation, participants sequentially viewed two encoding displays, each containing two colored oriented bars, separated by a delay. After a second delay, participants were probed to reproduce the orientation of one of the colored bars using a response dial. The handles of the response dial could be turned clockwise by pressing and holding the ‘M’ key with their right hand, and counterclockwise by pressing and holding the ‘Z’ key with their left hand. (**B**) Memory load manipulation. Preceding a block of trials, participants were instructed to selectively memorize the orientation of one, or two colored bars. In load-one, this bar could occur in either of the two encoding displays (early, or late). In load-two, each of the two cued bars were presented in a different encoding display (one early, the other late). (**C**) Manipulation of the tilt-response mapping. In addition to memory load, the starting position of the response dial was manipulated between blocks: it could be either vertical, or horizontal. Given that participants could only turn the dial handles a maximum of 90 degrees in either direction, the mapping between tilt and response varied throughout the experiment. Given a vertical response dial, a leftward tilted bar could only be reproduced using the left hand, and a rightward tilted bar using the right hand. Alternatively, given a horizontal response dial, a leftward tilted bar could only be reproduced using the right hand, and a rightward tilted bar using the left hand.

First, the orientation (left- or rightward tilt) of the bar was linked to the required response hand, and the mapping between tilt and response hand was counterbalanced by varying the starting point of response dial between blocks (**Figure 1C**). This uniquely enabled us to disambiguate prospection of the later required response from the visual tilt, since the same tilt was associated with different response hands in different blocks. Second, participants were instructed via a blocked pre-cue to selectively encode one (load-one), or two (load-two) colored oriented bars (**Figure 1B**). Crucially, in load-one, the memorized orientation was *certain* to become relevant for behavior during the memory delay, whereas in load-two, either of the two bars could *potentially* become relevant for behavior. This allowed us to test whether prospective responses with less certainty of relevance also evoked action encoding. Third, the cued oriented bars were sequentially presented to the participant in two different encoding displays (**Figure 1A**). This allowed us to isolate and track action planning for each potential action in load-two, and consider the role of this action planning on eventual behavior. We anticipated that planning for the potential action would be particularly clear in response to the first display, given that no prior action-plan could have been formed yet.

### Task performance depends on memory load and is comparable across tilt-response mappings

Before turning to our three main questions of interest (see introduction), we considered the effects of memory load (one/two), response dial (vertical/horizontal), and target moment (early/late) on task performance. We considered both decision times and errors. Decision time was defined as the time (in seconds) between probe onset and response initiation. Error (in degrees) was defined as the absolute difference between the orientation of the probed memory item and the reported orientation.

Participants were significantly slower (**Figure 2A**; *F*(1,192) = 98.92, *p* < 0.001, *η^2^* = 0.34) and less precise in their orientation-reproduction report in load-two, compared to load-one(**Figure 2B**; *F*(1,192) = 123.50, *p* < 0.001, *η^2^* = 0.37). Additionally, we observed that participants were more precise at reproducing the orientation when the probed target item was in the second encoding display (**Figure 2B**; *F*(1,192) = 12.29, *p* < 0.001, *η^2^* = 0.037). However, we did not observe such an effect for decision times (**Figure 2A**; *F*(1,192) = 0.034, *p* = 0.854, *η^2^* = 0.0001). Importantly, we observed no effect of response-dial type on either decision time (*F*(1,192) = 1.638, *p* = 0.202, *η^2^* = 0.006) or absolute error (*F*(1,192) = 0.621, *p* = 0.431, *η^2^* = 0.002), suggesting task difficulty was comparable across tilt-response mappings. We also observed no significant interactions between any of the variables on either measure of task performance (all *p* > 0.05).

**Figure 2.**
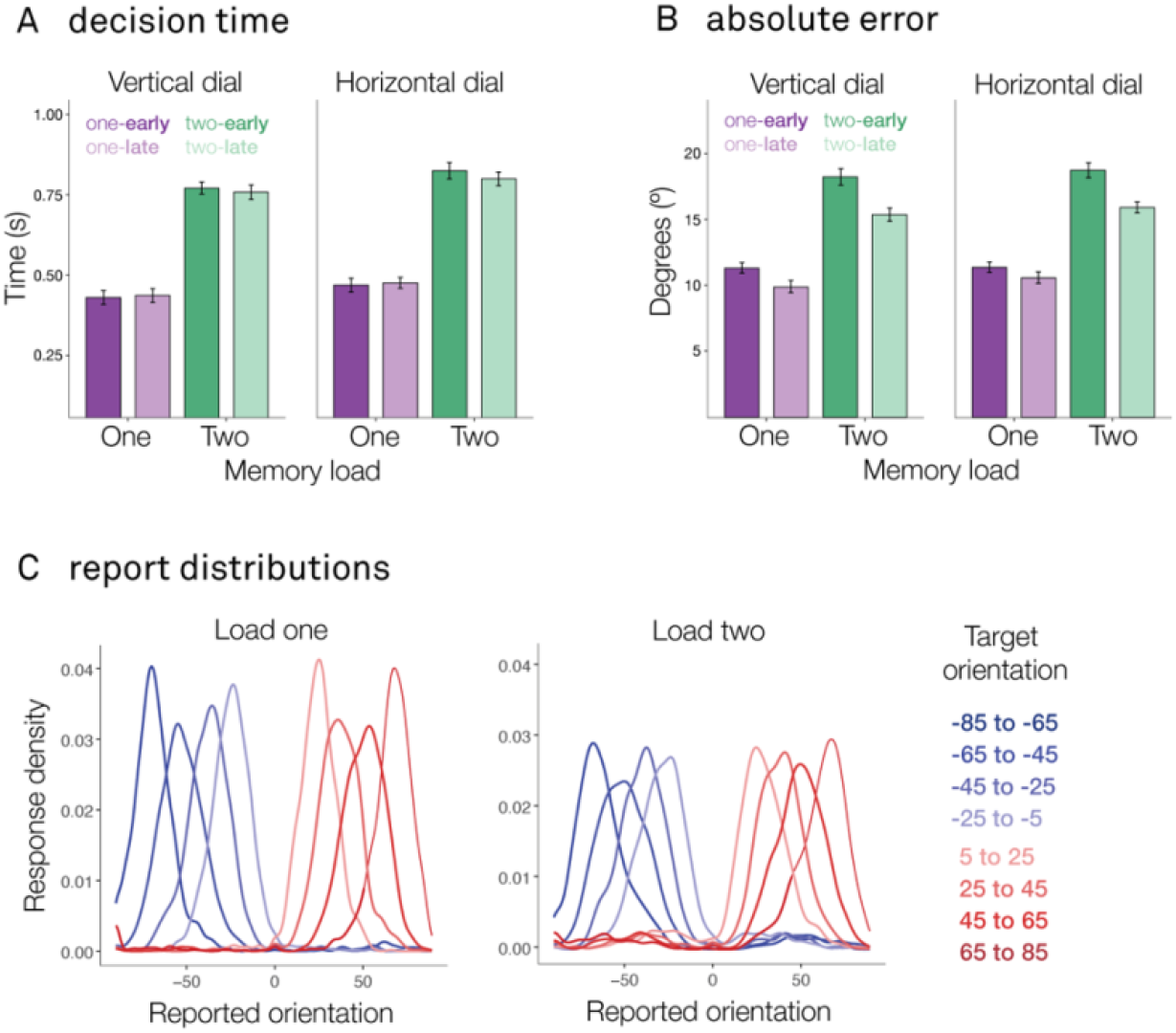
Task performance is comparable across tilt-response mappings. (**A, B**) Bar graphs show the average decision time (in seconds) and absolute error (in degrees) of the orientation-reproduction reports as a function of memory load (one/two), response-dial type (vertical/horizontal), and target moment (early/late). Error bars represent within-participant standard error (SE). (**C**) Density plot shows the distribution of the reported orientation as a function of the target orientation (in 20-degree bins), for load-one and load-two separately.

Finally, reported-orientation distributions for different target orientations (**Figure 2C**) revealed how participants guided their orientation-reproduction reports by the *precise* visual orientation of the probed memory item, rather than merely remembering whether to press the left or the right button at the end of the memory delay. This is apparent both in load-one (left panel) and in load-two (right panel), while also confirming the reduced overall precision in load-two (quantified above).

### Action encoding flexibly codes for the anticipated task

We now turn to our first main question: do we find action encoding even when the tilt-response mapping is varied across blocks (i.e., the same visual tilt is associated with different response hands in different blocks, as shown in **Figure 1C**)?

An established neural marker of manual-action planning in the EEG-signal is the attenuation of beta-band activity in electrodes contra-versus ipsilateral to the required response hand (Mcfarland et al., 2000; Neuper et al., 2006; Salmelin & Hari, 1994; Van Wijk et al., 2009), including in the context of prospective visual-working-memory-guided behavior (Boettcher et al., 2021; Ester & Weese, 2022; Nasrawi & van Ede, 2022; Rösner et al., 2022; Schneider et al., 2017). Here, we tracked this action-planning signal during the memory delay for each memory load condition (one/two). In load-one, we did so separately for early and late targets (occurring either in the first, or second encoding display). In load-two, the two potential targets were presented across the two encoding displays, and always required a different response hand. Accordingly, we could collapse over the early and late target trials, and defined contra-versus ipsilateral relative to the required response hand associated with the target-tilt in the first encoding display.

In line with previous studies (Boettcher et al., 2021; Nasrawi & van Ede, 2022; Schneider et al., 2017), we show a clear attenuation of beta-band power in motor electrodes contra-versus ipsilateral to the required response hand when the planned action is *certain* to become relevant (i.e., load-one; **Figure 3A-i**; early target T1, cluster p < 0.0001; late target T2, cluster p = 0.0001). As seen in the beta-band time-courses (**Figure 3A-ii**), these effects arise early after the onset of the relevant visual encoding display (early target T1, cluster p = 0.04; late target T2, cluster p = 0.04). This is in line with an early ‘encoding’ of the prospective action (as in Boettcher et al., 2021) that is certain to become relevant after the memory delay. Following this initial early post-encoding lateralization, we observed another clear attenuation of beta-band power preceding the memory probe, in anticipation of response execution (early target T1, cluster p < 0.0001; early target T2, target cluster p = 0.0001). Topographies, showing the difference between trials that required a left- or righthand response, confirmed that these motor signatures were particularly prominent in the corresponding left and right motor electrodes (C3/C4). This was the case both early after visual encoding, and before the probe-onset (**Figure 3A-ii**).

**Figure 3.**
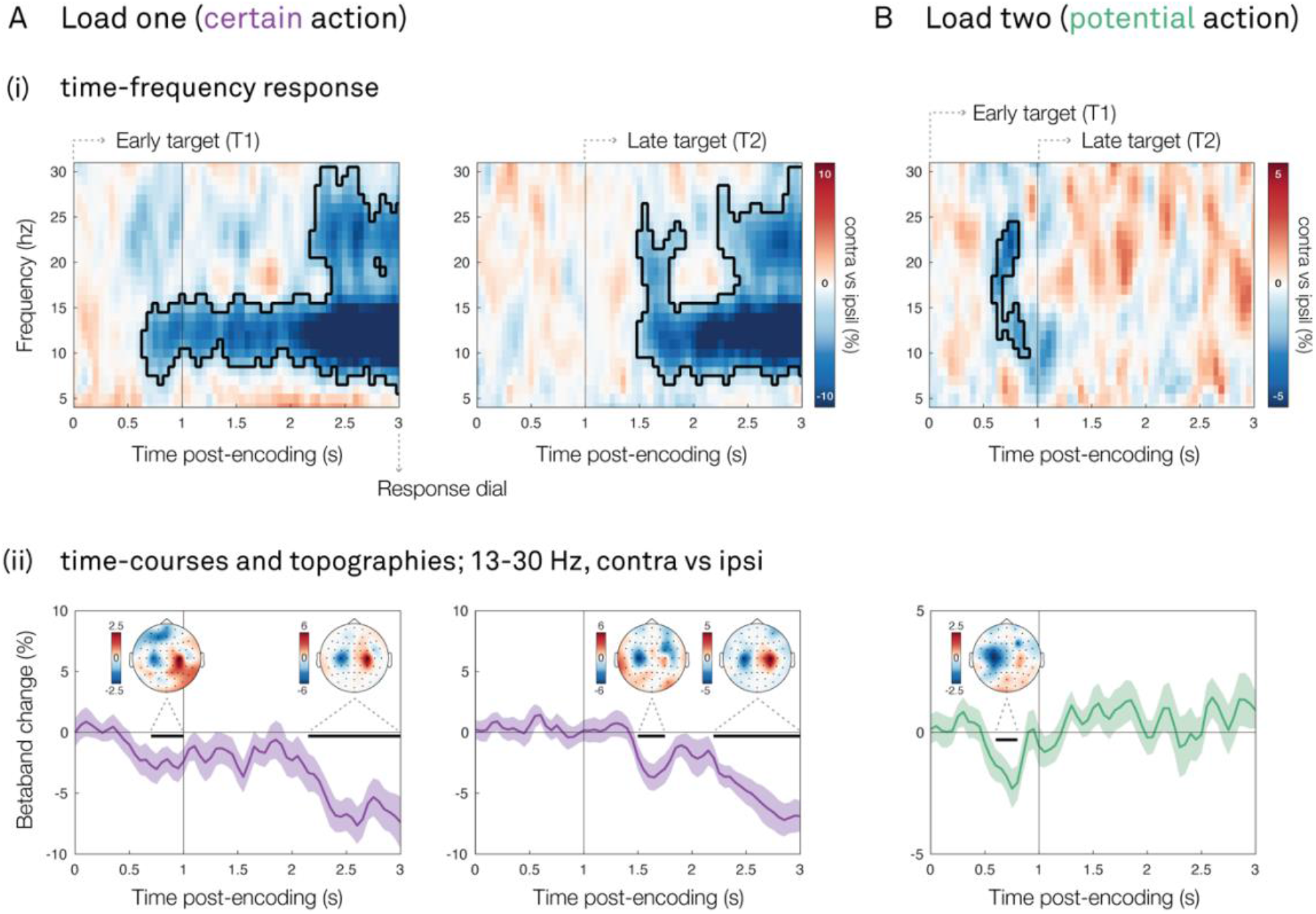
Action encoding flexibly codes for the anticipated task, and even when this is a potential task. For (**A**) load-one (certain action) and (**B**) load-two trials (potential action): (**i**) Lateralized motor activity relative to the response hand that is required to reproduce the visual item, calculated in canonical motor electrodes (C3/C4). The black outlines indicate significant clusters yielded from cluster-based permutation analysis. The vertical black lines indicate the onset of the encoding displays. (**ii**) Time-courses of lateralized motor activity at 13-30Hz (as in Boettcher et al., 2021). The black horizontal lines indicate significant clusters yielded from cluster-based permutation analysis. The vertical black lines indicate the onset of the encoding displays. Topographies show associated 13-30Hz motor lateralization (left vs. right required response hand), for each significant time-course-cluster. Shadings indicate standard error (SE) across participants.

The data from the load-one condition alone already provide an important advance over previous studies that linked visual features of memory content to manual actions (Boettcher et al., 2021; Ester & Weese, 2022; Henderson et al., 2022; Nasrawi & van Ede, 2022; Rösner et al., 2022; Schneider et al., 2017; van Ede et al., 2019). By varying the tilt-response mapping we could isolate action planning signatures from signatures associated with specific visual tilts because in half the blocks the same visual tilt would be associated with the opposite response hand (see Methods for details). Accordingly, our observed action encoding (**Figure 3A**) cannot be due to a fixed automatic mapping between visual tilt and a manual action. Instead, it must be attributed to flexible prospection of the upcoming task demand.

### Action encoding occurs for both certain and potential prospective actions

We now turn to our second main question: are, in addition to certain future actions, potential future actions also prospected when sequentially encoding multiple visual items?

To address this question, we turned to the data from our load-two condition. Strikingly, we observed a similar lateralization of beta-band power after visual encoding when the planned action could *potentially* become relevant, as was the case in load-two (**Figure 3B-i**; cluster p = 0.01). Here the chance that the encoded item would be probed for report was only 50%. Similar to what we observed in load-one, this signature was apparent early after the onset of the first visual encoding display (time-course cluster p < 0.02), and was again associated with a modulation in canonical motor electrodes (**Figure 3B-ii**).

Thus, in addition to certain actions, potential actions are encoded immediately after sensory encoding (referred to as *“output planning at the input stage”* in Boettcher et al., 2021). This suggests that the brain prospects a potential future manual action that is associated with a visual stimulus, even when this action *might* become relevant after the memory delay.

### Action encoding facilitates performance at the expense of subsequent memory items

Finally, we considered how potential-action encoding relates to subsequent memory-guided behavior at the end of the memory delay: does stronger encoding of a potential future action benefit subsequent behavior? And if so, does it come at the expense of performance for other memory items?

To test this, we zoomed in on the potential-action encoding effect in load-two (**Figure 3B-i**), where there were two items that competed in working memory. We averaged over the frequency-band (**Figure 4A**) and time-window (**Figure 4B**) of the observed potential-action encoding cluster. We then looked at beta lateralization in this time-frequency window (as a marker for the degree of action encoding) and compared trials with *fast* versus *slow* decision times, separated using a median split. Crucially, we did this separately for trials where the probed item was an *early* (T1) versus a *late* (T2) target.

**Figure 4.**
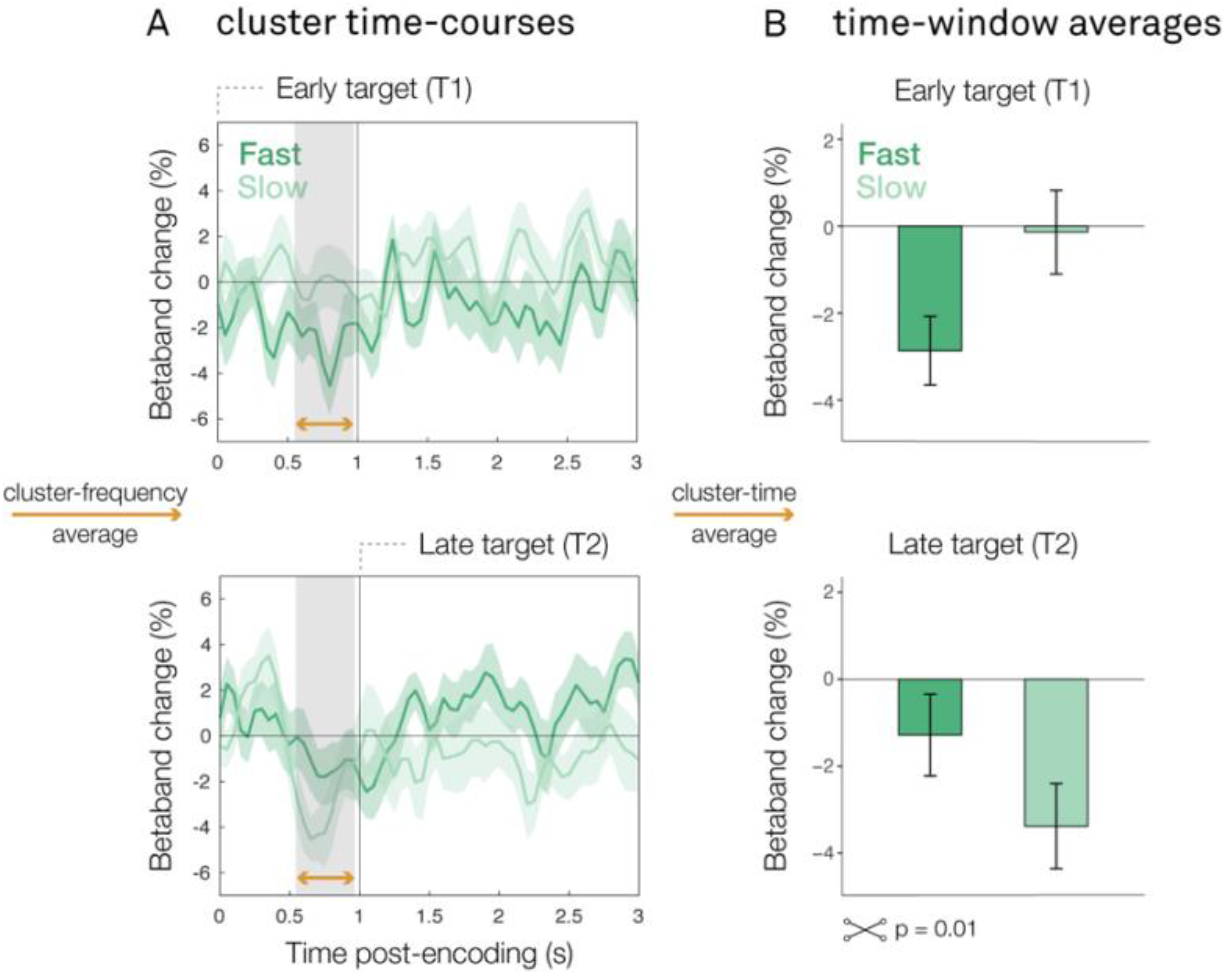
Stronger potential action encoding facilitates performance at the expense of subsequent memory items. **(A)** Time-course of the lateralized motor activity cluster observed in load-two, calculated in canonical motor electrodes (C3/C4). (**B**) Bar graphs of the lateralized motor activity cluster observed in load-two. Top panels show the response in trials where the target occurred in the first encoding display (T1, early). Lower panels show the response in trials where the target occurred in the second encoding display (T2, late). Dark green time-courses represent trials that ultimately had fast decision times; light green time-courses represent trials that ultimately had slow decision times; trials were marked as fast/slow using a median split. Error bars represent within-participant standard error (SE).

This revealed a significant interaction between the moment of the probed item (early/late target) and the speed of memory-guided behavior (fast/slow) regarding the degree of potential-action encoding after the first encoding display (*F*(1,96) =6.84, *p* = 0.01). The pattern of this interaction is consistent with a performance trade-off: when participants were probed to reproduce the orientation of the item in the first display (**Figure 4**, *top panels),*they were faster if they had more strongly encoded the action associated with this item (i.e., more beta lateralization in our action-encoding time-window of interest). However, if they were subsequently probed to reproduce the orientation of the item in second encoding display (**Figure 4**, *bottom panels),* participants responded *slower* after having more strongly encoded the potential action associated with the visual item in the first encoding display.

In other words, stronger encoding of a potential future action facilitates later memory-guided performance for the associated memory content, but this comes at the expense of responding to other potentially relevant and subsequently encoded working-memory content.

## DISCUSSION

It is increasingly appreciated that visual working memory does not solely serve to reflect on the past, but also to prepare for the future (Myers et al., 2017; Rainer et al., 1999; van Ede & Nobre, 2023). In line with this perspective, many recent studies have revealed close links between visual working memory and action (as reviewed in Heuer et al., 2020; Olivers & Roelfsema, 2020; van Ede, 2020). For instance, it has been shown that action planning takes place alongside retention of visual representations in working memory (Boettcher et al., 2021; Ester & Weese, 2022; Henderson et al., 2022; Kikumoto et al., 2022; Nasrawi & van Ede, 2022; Rösner et al., 2022; Schneider et al., 2017; Trentin et al., 2023; van Ede et al., 2019). Specifically, Boettcher et al. (2021) demonstrated that such output planning may already commence at the input stage, a pattern that was referred to as “action encoding”. The current study advances this growing body of research in three ways.

First, we show that action planning – as reflected by the pattern of action encoding during the memory delay – truly reflects the early prospection of a future required task, as it flexibly adapts to different visual-motor mappings. Similar to previous studies (Boettcher et al., 2021; Henderson et al., 2022; Rösner et al., 2022; Schneider et al., 2017; van Ede et al., 2019), we linked visual tilt (left- or rightward) to specific manual responses. Critically, however, in the current study we additionally varied the mapping between tilt and response hand, such that any specific tilt required reproduction with a left hand in some blocks, and with the right hand in other blocks. Despite this manipulation, we still observed clear action encoding when considering EEG motor signals contra-versus ipsilateral relative to the required response hand (and thus independent of the visual tilt). This shows that action encoding cannot reflect an automatic response to a specific visual orientation, but instead reflects a genuine prospection of the future task that is required to reproduce that orientation.

Second, we show that early action encoding occurs not only when this action is *certain* to become relevant (as in our load-one condition; and as previously shown by Boettcher et al., 2021), but also when this action is *potentially* relevant for future behavior. In everyday life, we are often required to encode and retain multiple visual items in working memory that each afford different actions, of which only some may become relevant for behavior depending on how the situation unfolds. Indeed, a key function of working memory may be to enable us to be ready for multiple potential courses of action (Cisek & Kalaska, 2005, 2010; Nasrawi & van Ede, 2022; van Ede et al., 2019; van Ede & Nobre, 2022). In the current study, we show that the observation of “output planning at the input stage” (Boettcher et al., 2021) extends to the planning of potential outputs: action encoding occurs even when participants know they will have to encode another item into working memory that is equally likely to become relevant for behavior after the memory delay. Thus, even when an action might only potentially become relevant in the future, it is still prospected immediately after visual encoding.

Third, by considering action encoding in a situation with multiple items in memory, we were in a unique position to reveal that stronger potential action encoding for one item may come at the expense of another. Specifically, we observed a performance trade-off, where stronger action encoding for one item is beneficial to performance when this item is subsequently probed, but hampers performance when the other item is probed. This reveals that there is variation in the degree to which a potential action is encoded – that is, potential actions may be more or less strongly prioritized – which is reflected in performance later on. This is consistent with recent research showing that actions that were made or planned during visual working memory can affect performance of action-matching items, at the expense of other items (Hanning et al., 2016; Hanning & Deubel, 2018; Heuer & Schubö, 2017, 2018; Ohl & Rolfs, 2017, 2018, 2020; Trentin et al., 2023).

In our load-one condition, we observed action encoding both when the relevant item occurred in the first and second display (see **Figure 3**). Interestingly, however, in our load-two condition, we only observed action encoding following the first display, with little evidence for a similar effect following the second display. Different factors may account for this. In our design, the relevant items in the first and second display were always linked to different response hands. In our analysis, we focused on lateralization of beta-band power relative to the required response hand in the first display. After the first encoding display, only one potential action could be planned (reflected in the observed ‘potential action encoding’ effect). In contrast, after the second display, participants were required to add a second potential action, while maintaining the action associated with the first display. Accordingly, the lack of clear potential action encoding following the second display may be due to both potential actions balancing out relative beta-band change, as they each require a different response hand. Alternatively, the first potential action may be selectively planned, while the second one is ignored. However, this alternative explanation would predict a clear decision-time benefit for the item in the first display, but we found no evidence for this (see **Figure 2**).

In our paradigm, the encoding of multiple visual items into working memory occurred sequentially. This manipulation allowed us to track action encoding for each memory item separately, advancing previous research where multiple visual items were presented at the same time (Nasrawi & van Ede, 2022; van Ede et al., 2019). Yet, together with the advance of this manipulation, we also lost the possibility to infer whether multiple potential actions were planned and encoded in parallel. From dedicated action-planning studies, there is evidence that multiple actions can be planned in parallel, before selecting the relevant action for implementation (Cisek, 2007; Cisek & Kalaska, 2005; Gallivan et al., 2015, 2016; Grent-’t-Jong et al., 2013; but cf. Dekleva et al., 2018). Whether this also holds for the encoding of multiple potential actions (i.e., parallel action encoding) in the context of visual working memory tasks remains an exciting avenue for future research.

In summary, our findings confirm that action planning – and specifically action encoding – alongside visual working memory reflects anticipated task demands, rather than an automatic response to visual tilt; occurs even for actions that will potentially become relevant for behavior; and is associated with better performance, at the expense of performance to other memory content. Together, these results demonstrate that the potential prospective use of visual working memory content is flexibly planned for early on, with consequences for later performance.

## METHODS

### Participants

Twenty-five healthy human adults (age: mean = 22.85, sd = 4.03; gender: 17 female, 8 male; 4 left-handed) participated in the experiment. All participants had normal, or corrected-to-normal vision, and none were excluded from the analyses. The experiment was approved for by the Research Ethics Committee of the Vrije Universiteit Amsterdam. Participants provided written informed consent before participating in the study, and were rewarded €10 or 10 research credits per hour for their participation.

### Experimental design and procedure

The experiment was programmed in PsychoPy (Peirce et al., 2019). All stimuli were presented on a led monitor (ASUS ROG STRIX XG248; 23.8 inch, 1920×1080p, 240Hz) situated 60 cm away from the participant. Each trial contained two separate visual encoding displays (**Figure 1A**), with two colored oriented bars presented on either side of a fixation cross, at 4° visual angle distance from fixation. One bar was always tilted to the left, the other to the right, and could be presented either to the left or right of the fixation cross. The location of the left- and rightward oriented bar remained fixed between the two encoding displays during a trial. The magnitude of the orientation of the bars was randomly determined, and varied between 5° and 85° to avoid fully vertical, or fully horizontal orientations. Each bar had a width of 0.4° and height of 4° visual angle, and could have one out of four possible colors: yellow (HEX-value: #C2A025), blue (#3843C2), green (#2FC259), or red (#CF3C3C).

After the second working-memory delay, the fixation cross changed to the color of one the oriented bars, probing participants to reproduce the orientation of the color-matching bar using a response dial. The response dial, presented around fixation, consisted of a large circle with the same diameter as the bar-length with two smaller circles (‘handles’) placed on opposite points on the circle, representing an orientation. The position of the response dial handles could be adjusted with a counterclockwise or clockwise turn (max 90° in each direction), by pressing and holding the ‘Z’ or ‘M’ key respectively. Participants could only press one key per response, and the dial would move clock-wise (Z) or counter-clockwise (M) for as long as the button was pressed. Participants finalized their orientation report by releasing the chosen key when hitting the desired orientation. Immediately after report finalization, participants received feedback on their performance. This feedback consisted of a number between 0 and 100%, representing the percentage of overlap between the reported- and target orientation.

We linked visual tilt to the required response hand as follows: a left- or rightward tilted bar could only be accurately reported by pressing and holding a key with either the left or right hand. Extending our prior studies that leveraged this manipulation (Boettcher et al., 2021; van Ede et al., 2019), we here used two different tilt-response mappings (**Figure 1C**) that varied across blocks. When the start position of the response dial was vertical (0°), a leftward tilt (relative to the vertical starting point) could only be accurately reported with the left response hand (an ‘Z’ key), and a rightward tilt with the right response hand (an ‘M’ key). When the start position of the response dial was horizontal (90°), a leftward tilt could only be accurately reported with the right response hand, and a rightward tilt with the left response hand (see examples in **Figure 1C**).

Preceding each block of trials, participants were presented with a pre-cue informing them of the color(s) of the relevant bar(s) that block (**Figure 1B**). In *load-one* blocks, participants were instructed to memorize the orientation of a single bar (e.g., only the blue bars’ orientation), which was *certain* to be probed for reproduction report later. In *load-two* blocks, participants were instructed to memorize the orientations of two bars (e.g., both the yellow and red bars’ orientation), which could each *potentially* be probed for reproduction report (randomly selected, and known to the participant after the memory delay). In both load conditions, target (i.e., the probed item) moment (first, or second encoding display), target location (left, or right side of the screen), and target tilt (left, or right) were counterbalanced within a block. In load two, the non-target (i.e., the non-probed item) was always present at a different moment, at a different location, and of a different tilt-type.

Preceding the main experiment, participants practiced the task for 5-10 minutes, or until their performance was good enough (~75% precision of their orientation-reproduction report). They then completed two consecutive sessions of approximately 45 minutes, with a self-paced break in between. Each session consisted of 12 blocks of 32 trials (384 trials per session). Response dial and memory load type were blocked and counterbalanced, and occurred in a random sequence. The response dial type always remained the same for a minimum of 64 trials. Each dial type block was preceded by 4 dial-practice trials in which participants practiced turning the response dial clockwise and counterclockwise to match a visible orientation (no working memory component). This was done to avoid confusion about the response dial type during the main trials. These trials were not included in the presented analysis.

### Behavioral analyses

Behavioral data were analyzed using R (R Core Team, 2020). Absolute orientation-reproduction error (in degrees) was defined as the absolute difference in orientation between the target and report. Decision time (in seconds) was defined as the time between probe onset and response initiation. Trials with decision times lower than 100 msec or larger than 5 sec were excluded from further analyses. In addition, for each participant, trials were excluded with decision times larger than the mean plus 2.5 times the standard deviation (this was the case for less than ~3% of the trials). Two one-way repeated measures ANOVAs were performed to evaluate the effect of memory load (one/two), target moment (early/late), and response dial (horizontal/vertical) on the absolute error and decision times. All effects were visualized using the ggplot2 package (Wickham, 2016).

### EEG acquisition and analyses

#### Acquisition

EEG was measured using the BioSemi ActiveTwo System (biosemi.com) with a standard 10–10 System 64 electrode setup. Two electrodes were placed on the left and right mastoid, and used for offline re-referencing of the data. Additionally, EOG was measured using one electrode next to, and another above the left eye.

#### Preprocessing

All EEG analyses were performed in MATLAB (2022a; The MathWorks, 2020) using the FieldTrip toolbox (Oostenveld et al., 2011; https://fieldtriptoolbox.org). Data were epoched from −1000 to 4000 msec, relative to the onset of the first encoding display, and re-referenced to an average of the left and right mastoids. Noisy channels (if present) were interpolated by taking the average of two adjacent electrodes. Next, the data were down-sampled to 200Hz. Independent component analysis (ICA) was used to correct for blink artifacts. The appropriate ICA components used for artifact rejection were identified by correlating the time-courses of the ICA components with those of the measured horizontal and vertical EOG. After blink correction, the FieldTrip function *ft_rejectvisual* was used to visually assess which trials had exceptionally high variance, which were marked for rejection. Trials that had been marked as too fast or too slow (as described in the Behavioral Data Analyses section) were also rejected from further analyses. A surface Laplacian transform was applied to increase the spatial resolution of the central motor beta signal of interest (as also done in Boettcher et al., 2021; Nasrawi & van Ede, 2022; van Ede et al., 2019).

#### Electrodes, frequency-band, and time-window selection

For all analyses, channel and frequency-band selections were set a priori, and in line with previous research (Boettcher et al., 2021; Mcfarland et al., 2000; Neuper et al., 2006; Salmelin & Hari, 1994; Van Wijk et al., 2009). To investigate motor planning signals contra-versus ipsilateral to the required response hand, we focused on activity in EEG electrodes C3 and C4. Additionally, we extracted activity in the beta-band frequency (13-30Hz) for all our time-course visualization.

#### Time-frequency analysis

Time-frequency responses were obtained using a short-time Fourier transform of Hanning-tapered data. A 300 msec sliding time window was used to estimate spectral power between 3 and 40 Hz (in steps of 1 Hz), progressing in steps of 50 msec. Next, activity in motor electrodes (C3/C4) was contrasted between trials in which the required response hand was contralateral versus ipsilateral to these electrodes. This was expressed as a normalized difference: ((contra-ipsi)/(contra+ipsi)) x 100. These contrasts were then averaged across the left and right motor electrodes. To extract time-courses of lateralized motor activity, we averaged this contra-versus ipsilateral response across the 13-30Hz beta band. Topographies of the lateralized motor activity were obtained by contrasting trials in which the memory content was associated with a left vs. right hand response, expressed as a normalized difference between left and right trials, for each electrode.

#### Relation between EEG action-encoding and subsequent behavior

We asked whether the degree of potential-action encoding during the memory delay varied depending on subsequent behavior. For this analysis, we focused on load-two trials where we could address whether there may be a trade-off whereby stronger action encoding for one item may benefit performance for that item at the expense of another. To test this, we defined an action-encoding window of interest in terms of frequency and time bounds based on the action-encoding cluster previously observed in load two. Importantly, as this cluster was obtained from all load-two trials, the selection of this window of interest was made irrespective of subsequent behavior and target moment. For these extracted data of interest, we then compared trials with fast vs. slow decision times after the memory delay (based on a median split), to see whether the pattern of action encoding after visual encoding could predict the speed of onset of subsequent working-memory guided behavior (as in Boettcher et al., 2021). Here, we did this separately for trials where the probed item occurred in the first or second display. A one-way repeated measures ANOVA was used to evaluate how the action-encoding differed depending on decision times (fast/slow) and target moment (early/late). These effects were visualized using the ggplot2 package (Wickham, 2016).

#### Statistical evaluation

Cluster-based permutations (Maris & Oostenveld, 2007) were performed for the statistical evaluation of the above-described EEG contrasts. This nonparametric approach (or Monte Carlo method) offers a solution for the multiple-comparisons problem in the statistical evaluation of EEG data, which, in our case, included a sizeable number of time–frequency comparisons. It does so by reducing the data to a single metric (e.g., the largest cluster of neighboring data points that exceed a certain threshold) and evaluating this (in the full data space under consideration) against a single randomly permuted empirical null distribution. Cluster-based permutations were performed on the time–frequency responses (considering clusters in time and frequency) and time-courses (considering clusters in time) using 10.000 permutations, and an alpha level of 0.025.

## Acknowledgements

This research was supported by an ERC Starting Grant from the European Research Council (MEMTICIPATION, 850636) and an NWO Vidi grant by the Dutch Research Council (grant number 14721) to F.v.E. The authors thank Babak Chawoush and Merlijn Breunesse for their valuable comments on the manuscript.

## Author contributions

R.N. and F.v.E. designed and conceptualized the research; R.N. programmed the experiment, collected the data, and performed the analyses; R.N. wrote the original draft, and made the figures; R.N., S.E.P.B., and F.v.E. revised and edited the manuscript.

## Code and data availability

The <u>experiment code</u> and <u>analysis code</u> are both publicly available on GitHub. The data will be made available upon acceptance for publication.

